# Engineering bacteriophages through deep mining of metagenomic motifs

**DOI:** 10.1101/2023.02.07.527309

**Authors:** Phil Huss, Kristopher Kieft, Anthony Meger, Kyle Nishikawa, Karthik Anantharaman, Srivatsan Raman

## Abstract

Bacteriophages can adapt to new hosts by altering sequence motifs through recombination or convergent evolution. Where such motifs exist and what fitness advantage they confer remains largely unknown. We report a new method, Metagenomic Sequence Informed Functional Scoring (Meta-SIFT), to discover sequence motifs in metagenomic datasets that can be used to engineer phage activity. Meta-SIFT uses experimental deep mutational scanning data to create sequence profiles to enable deep mining of metagenomes for functional motifs which are otherwise invisible to searches. We experimentally tested over 17,000 Meta-SIFT derived sequence motifs in the receptor-binding protein of the T7 phage. The screen revealed thousands of T7 variants with novel host specificity with functional motifs sourced from distant families. Position, substitution and location preferences dictated specificity across a panel of 20 hosts and conditions. To demonstrate therapeutic utility, we engineered active T7 variants against foodborne pathogen *E. coli* O121. Meta-SIFT is a powerful tool to unlock the functional potential encoded in phage metagenomes to engineer bacteriophages.

## Introduction

Bacteriophages are powerful tools for targeting bacterial pathogens and making precise modifications to complex microbial communities. Phage genomes can be engineered to increase infectivity on different bacterial hosts or across different conditions, and even single mutations have been shown to improve phage activity over 1000-fold [1–3]. Despite this potential, high throughput exploration of complex, multiple-mutation alterations to phage genomes have largely remained elusive. A key challenge that must be overcome is determining which mutations are effective within the enormous sequence space that can be engineered in phages, as even small stretches of sequences present an overwhelming number of possible combinations. For example, a stretch of only six amino acids contains 6.4×10^7^ possible combinations of mutations, while a ten amino acid sequence has 1×10^13^ possible combinations. The number of possible combinations of mutations quickly exceeds our ability to create and assess phage variants through traditional passage of natural phages or by simple randomization. Effectively querying this immense sequence space to determine which sequences are most relevant to improve phage activity on targeted bacterial strains and understanding where we should incorporate these changes in the phage genome are key questions in phage engineering.

Here, we sought to answer this question using a novel approach leveraging phage metagenomic data to find sequence motifs that can drive phage activity. Metagenomic phage sequences are highly diverse and have been increasingly characterized, as demonstrated by the growing volume of viral metagenomes curated from the gut [4–7], oceans [8–11], lakes and soil microbiomes [12–15]. However, the function of many sequences remains largely unknown, and the general lack of sequence conservation among phages makes identifying functionally related metagenomic sequences difficult [9,16]. Even amongst closely-related phages, genes that are under constant evolutionary pressure can have little sequence homology [17]. For such highly divergent genes, we reasoned that sequence similarity could occur over short sequence motifs instead of full-length genes or domains. Convergent evolution may drive phages towards the same solution for binding to a particular receptor using a small motif in otherwise non-homologous sequences. However, the extent to which such motifs exist in metagenomic databases and if they can be harnessed to confer a fitness advantage to phages remains largely unknown.

In this study, we report a new method called Meta-SIFT (**<u>Meta</u>**genomic **<u>S</u>**equence **I**nformed **<u>F</u>**unctional **<u>T</u>**raining) to discover sequence motifs in metagenomic datasets that can alter phage activity. In contrast to exclusively bioinformatics-driven motif discovery tools, Meta-SIFT uses experimental deep mutational scanning (DMS) data to quantitatively score the importance of every possible substitution within a defined sequence window, preserving functionally important mutations, their relevant position, and curating motifs that can influence phage activity. This weighted substitution profile sifts through metagenomic databases to find relevant sequence matches. We used Meta-SIFT to curate viral metagenomic sequence motifs from the NCBI and IMG/VR databases for the tip domain of the receptor binding protein (RBP) of T7 phage. Because they are crucial determinants of phage activity, RBPs have proven excellent targets for engineering phage activity [1,2,18–25]. Out of a total sequence space of over 10^14^ possible combinations of 6mer or 10mer motifs at any position in the RBP tip domain, Meta-SIFT refined 10^4^ motifs from diverse, non-homologous metagenomic phage sequences to incorporate into the tip domain of the T7 RBP.

We performed pooled screening of T7 phages engineered to contain Meta-SIFT motifs against a panel of twenty bacterial hosts or environmental conditions, revealing thousands of phage variants with novel host activity and specificity. These motifs were well distributed in metagenomic space, indicating that deep mining of metagenomic datasets is a powerful tool for tailoring phage activity. Position and substitution preference across different hosts showed key combinations of substitutions that drive activity and host range. To demonstrate the therapeutic utility of this approach, we used Meta-SIFT to identify variants that were effective on foodborne pathogen *E. coli* O121 in conditions where neither wild type nor a single site saturation library has any activity. These results showcase the utility of Meta-SIFT to cross sequence space to find complex, high value combinations of mutations that improve phage activity on relevant pathogens that are otherwise unattainable through evolutionary selection.

## Results

### Meta-SIFT identifies diverse motifs from metagenomic phage sequences

Three considerations guided the development of our approach. First, relevant sequence motifs may be hidden in metagenomic proteins with little sequence homology to the original phage protein. A homology search using BLAST of the T7 RBP tip domain identified 83 unique sequences, an insignificant fraction of the potential matches in metagenomic sequences. Second, metagenomic motifs with a pronounced impact on phage activity may bear little resemblance to the wildtype motif. Thus, we needed to create a profile of query motifs instead of just the native sequence. Third, since motif activity depends on local protein context, we needed to determine where motifs needed to be inserted to best influence phage function.

We developed Meta-SIFT (**<u>Meta</u>**genomic **<u>S</u>**equence **I**nformed **<u>F</u>**unctional **<u>T</u>**raining), a bioinformatic motif-finding tool that uses experimental data from deep mutational scanning (DMS) to identify metagenomic motifs that influence phage activity (Figure 1A). Meta-SIFT is not constrained by overall sequence homology and can reach deep into metagenomic space for motifs from phages that are not closely related to the wildtype phage. Using DMS data to guide the search, Meta-SIFT evaluates every possible substitution at every position in candidate motifs, resulting in functionally relevant motifs that can significantly differ from the wildtype sequence. Using positional information from the DMS screen, Meta-SIFT also precisely identifies where motifs must be inserted within the phage to influence phage activity. As proof of concept, we used Meta-SIFT to mine metagenomic motifs for the RBP tip domain of the T7 phage.

**Figure 1.**
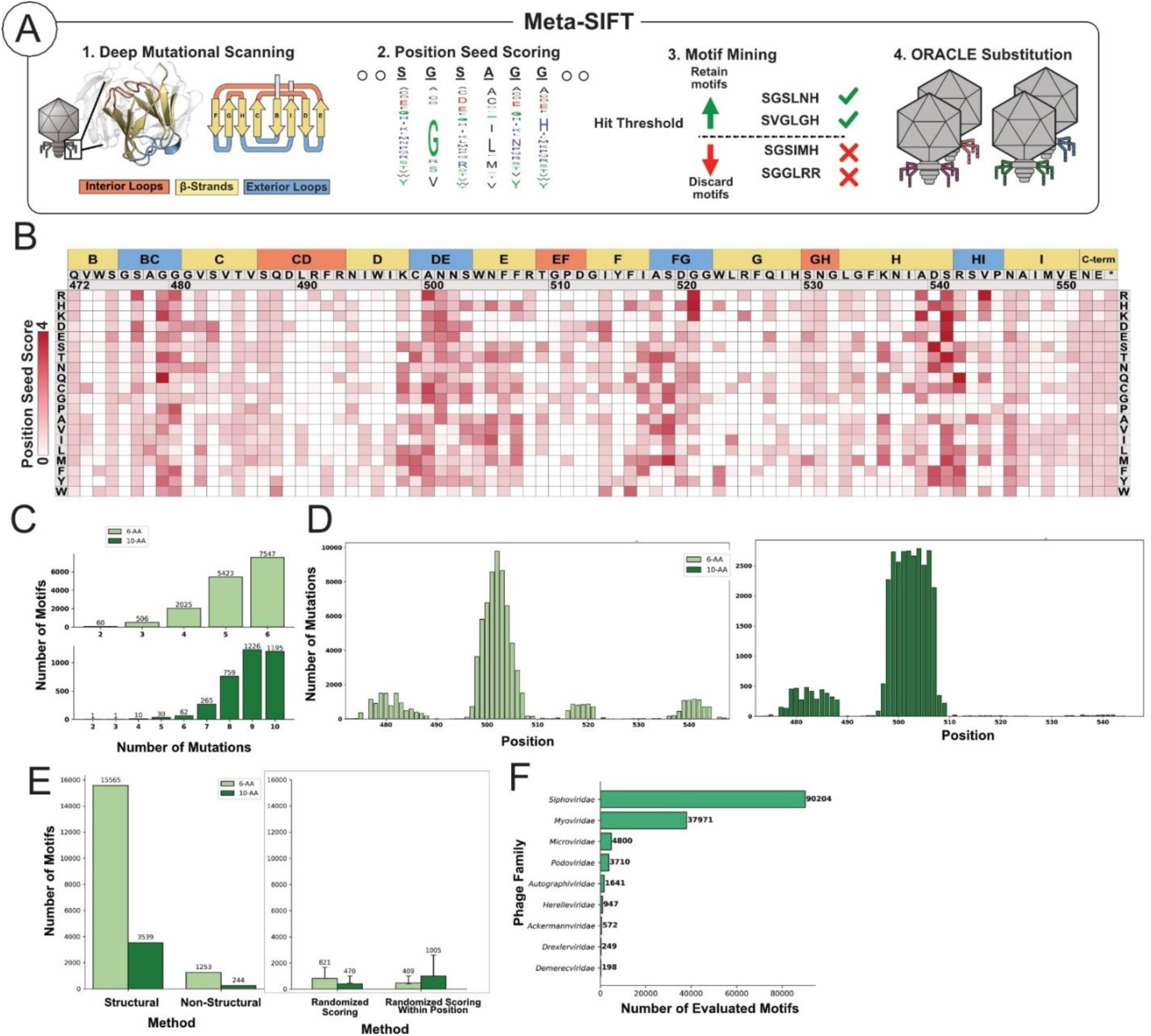
Meta-SIFT identifies diverse motifs from metagenomic phage sequences. **(A)** Illustration of Meta-SIFT. Results from DMS of the T7 phage RBP tip domain (1) are used to seed scores for probable motifs (2), which are then mined in metagenomic databases (3). Motifs passing a hit threshold are substituted into phages in a phage library using ORACLE (4). Crystal structure (PDB: 4A0T) and secondary structure topology shown color coded with interior loops as red, β-sheets as yellow and exterior loops as blue. **(B)** Heat map showing seed score (red gradient) for substitutions in the tip domain. Substitutions are shown top to bottom while position (residue numbering based on PDB 4A0T), wildtype amino acid and secondary structure topology is shown left to right. **(C)** Number of mutations per sourced motif for 6-AA (light green, top) or 10-AA (dark green, bottom) motifs. **(D)** The number of and relative position on the RBP of each mutation for 6mer (top, light green) and 10mer (bottom, dark green) motifs derived from the metagenomic dataset. **(E)** Number of motifs found passing hit threshold using Meta-SIFT and either metagenomic phage structural proteins or non-structural proteins (left). Number of motifs found using Meta-SIFT with metagenomic structural proteins after fully randomizing seed scores or randomizing seed scores at each position (right, mean ± SD of triplicate randomizations). 6-AA Motifs are shown in light green, 10-AA motifs shown in dark green. **(F)** Number of evaluated motifs sourced from different phage families derived from proteins annotated in NCBI.

Meta-SIFT first determines a ‘seed score’ for every possible substitution at every position using DMS data (Figure 1B, Supplementary File 1). We computed seed scores using a prior DMS study where we screened 1660 single-amino acid substitution variants of the T7 RBP tip domain on multiple hosts [2]. The seed score is generated based on the performance of each substitution at each position across evaluated hosts (see Methods). The seed score increases if that substitution resulted in a change in host range, indicating that position or kind of substitution is pivotal for changing phage activity. We imposed no limit on the number of mutations in any motif. This approach presents a notable tradeoff, as it predisposes motifs towards sequences that are expected to have poor activity on some bacterial hosts to improve the likelihood of identifying sequences that can drive distinct changes in host range. Seed scores are well distributed throughout the tip domain and favor positions known to influence phage activity such as exterior loops and outward-facing β-sheets (Figure 1B).

After compiling a list of candidate motifs derived from the seed scores, Meta-SIFT mines metagenomic datasets for matching sequences. We searched for motifs in publicly available NCBI and IMG/VR databases, representing some of the largest databases for metagenomic phage sequences [26,27]. We restricted our search to 6-mer and 10-mer motifs for simplicity and following the hypothesis that changes in phage activity can be influenced by short sequences. We ultimately mined motifs from ∼60,000 unique metagenomic phage structural proteins. We hypothesized productive motifs are likely to be evolutionarily enriched and retained only those motifs that met a minimum threshold of abundance. Our search yielded 15,561 6-mer and 3,549 10-mer protein motifs (Supplementary File 2). Mined motifs were highly dissimilar to the wildtype sequence, averaging 5.28±0.84 for 6mer motifs and 8.87±1.1 for 10mer motifs (Figure 1C). 6mer motifs were more well distributed than 10mer motifs, and the majority of both 6mer and 10mer motifs were located in and around the DE exterior loop and E β-sheet region (Figure 1D).

We used two approaches to evaluate if Meta-SIFT had identified motifs that were likely relevant to receptor recognition. First, we repeated Meta-SIFT using a comparably sized set of metagenomic non-structural proteins such as polymerases, ligases, and terminases, reasoning that fewer motifs for receptor recognition should be found in non-structural proteins. Indeed, Meta-SIFT curated ten-fold fewer motifs in non-structural proteins (Figure 1E Left). Second, we evaluated the importance of the DMS data by searching with randomized seed scores. Randomizing seed scores should result in fewer hits because scores lose relevance to phage activity, resulting in poorly seeded motifs that should be found less frequently than correctly seeded motifs. Randomizing either position or identity of the substitution scores (Supplementary File 1) produced ten-fold fewer motifs than correctly seeded scores, indicating our approach was finding appropriate motifs and emphasizing the importance of the DMS dataset for seeding scores (Figure 1E Right).

Finally, we evaluated if our approach sampled motifs from different phage families or was biased toward the *Podoviridae* family to which T7 phage belongs. We examined annotated protein taxonomy from source proteins from the NCBI database and found motifs were derived from diverse phages that mirrored the overall taxonomic distribution of the database, predominantly from the families *Siphoviridae* and *Myoviridae* (Figure 1F). Overall, these results suggest that Meta-SIFT can derive motifs that are distributed across diverse phage families using systematic experimental calibration of residue substitutions to guide the search.

### Meta-SIFT motifs direct activity across bacterial hosts

To evaluate how META-SIFT-derived motifs altered activity in the phage we synthesized all 19,109 candidates (15,561 6-mer and 3,549 10-mer protein motifs) as chip oligonucleotides with motifs inserted at the appropriate location in the tip domain (Supplementary File 2). The variant library was inserted into the T7 phage genome using ORACLE (**O**ptimized **R**ecombination, **Ac**cumulation, and **L**ibrary **E**xpression) [2]. ORACLE is a high-throughput precision phage genome engineering tool designed to create a large, unbiased library of phage variants of defined sequence composition. Variant libraries created using ORACLE are not required to be active on the host used to generate the library, and therefore the method is ideal for evaluating highly divergent sequences like the motif library that are likely to impose significant selective pressure. After chip synthesis and ORACLE engineering, 17,636 variants were detectable at sufficient abundance in the final phage library (92.3%, see Supplementary File 2). Following ORACLE, the expressed phage library retained a near-identical distribution of mutations, averaging 5.3±0.83 mutations for 6mer motifs and 8.88±1.09 mutations for 10mer motifs (see Figure S1A), with a similar distribution across the tip domain (see Figure S1B) and a normally distributed abundance with a small tail at greater abundance (see S1C), altogether indicating the library was successfully cloned and reflected the distribution of the original library.

We evaluated the activity of phage variants by selecting the library on a panel of twenty *E. coli* hosts or conditions that represented a range of different receptor profiles and susceptibility to wildtype T7 phage (Figure 2A-B). This panel included susceptible B strain derivative BL21, K-12 derivative BW25113, and DH10B derivative 10G, three uropathogenic strains of *E. coli*, seven different knockout variants of *E. coli* BW25113 with truncations to the phage’s lipopolysaccharide receptor, two knockouts of transporters *tonA* and *tonB* not expected to influence phage fitness, and *E. coli* O121, which was evaluated using 0% to 2% NaCl supplement representing the diverse range of salt conditions T7 phage can experience in the environment or in the human microbiome [2,28–33]. We were particularly interested in determining if Meta-SIFT could identify active variants of T7 on *E. coli* O121 in higher salt concentrations. Neither the wildtype nor a single amino-acid DMS library of the tip domain has any apparent activity in liquid culture or ability to plaque on this strain in this condition, indicating a solution to improve activity is too far in sequence space from the wildtype sequence to be present in the natural diversity of the phage population (Figure S1D).

**Figure 2.**
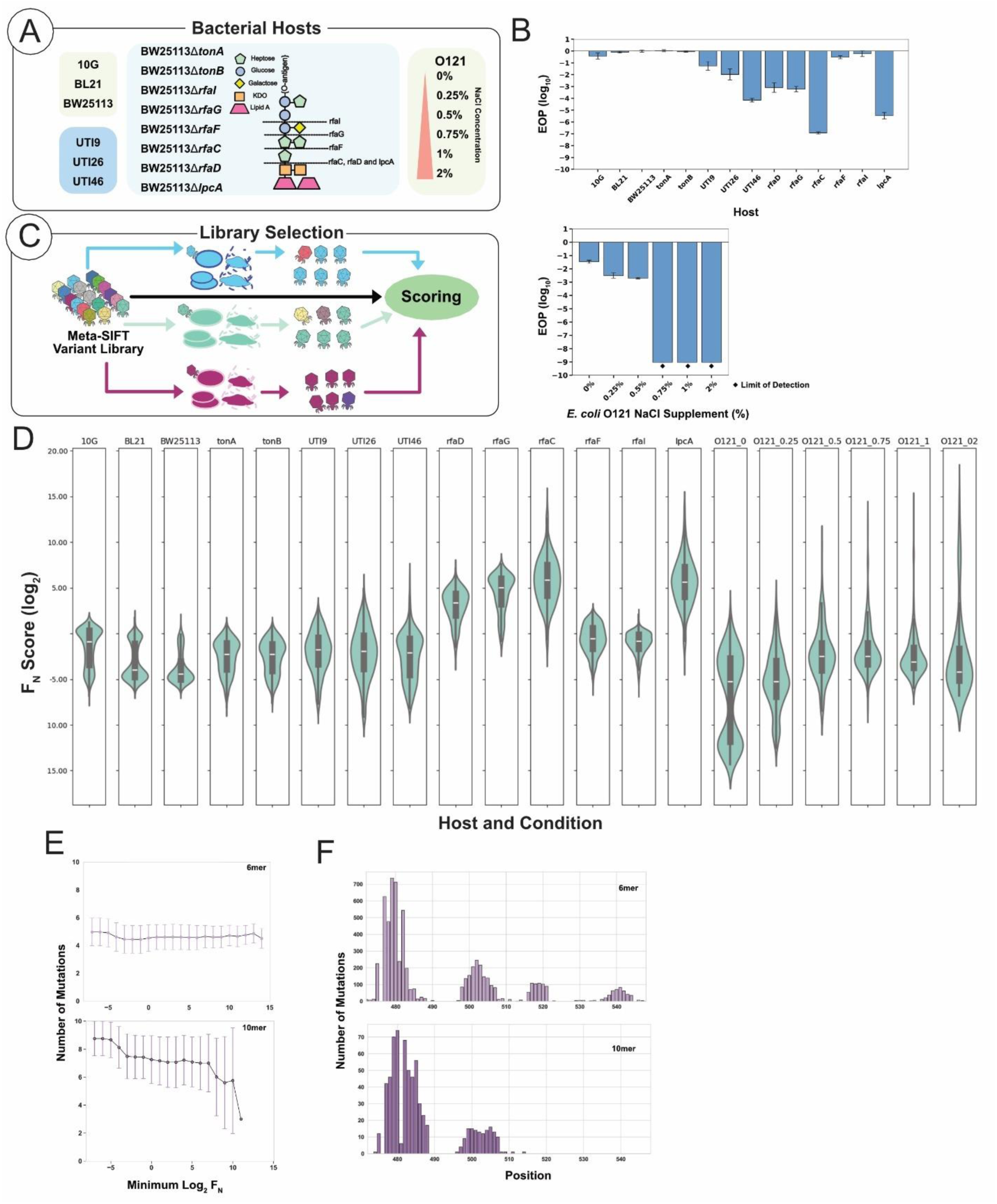
Motifs direct specificity and activity across bacterial hosts. **(A)** Schematic representation of evaluated host panel. Lipopolysaccharide structure for evaluated BW25113 bacterial hosts shown with the expected effect of the gene deletion on LPS structure denoted by a dashed line. **(B)** Efficiency of Plating (EOP) results shown for wildtype T7 phage on all evaluated hosts (top panel, BW25113 knockouts labeled as missing gene) and NaCl gradient for *E. coli* O121 (bottom panel), normalized to *E. coli* 10G (biological triplicates, +/- SD). 0.75%, 1% and 2% NaCl supplement reached the limit of detection for the assay with no apparent plaques. **(C)** Schematic representation of passage and scoring scheme. Library variants are scored by comparing variant abundance of an unbiased library pre- and post-selection on different hosts. NaCl concentration for O121 **(D)** Violin plots displaying the range of log_2_ F_N_ score for each evaluated host. BW25113 deletion mutants are labeled by their gene deletion, and O121 mutants by the percentage NaCl condition. **(E)** The average number of mutations (+/- SD) in each variant for 6mer (top, light purple) and 10mer (bottom, dark purple) motifs as activity (maximum F_N_ on any host or condition) increases. **(F)** The number of and relative position of each mutation for 6mer (top, light purple) and 10mer (bottom, dark purple) active motifs with an F_N_ > -3 after selection.

The library was selected on each host for approximately two replication cycles, after which the library was sequenced and variants assigned a functional score, F_N_, based on the ratio of the abundance of the variant before and after selection normalized to wildtype (Figure 2C). F_N_ scores were well correlated across biological duplicates (Figure S1F). In total 4,328 variants in the library (24.5% of the expressed library after ORACLE, Supplementary File 3) showed detectable activity on any one host, a high number given the number of impactful mutations present in each variant. The library displayed a dramatic range of activity on each host and contained high F_N_ variant that indicated Meta-SIFT successfully built phage variants that could improve activity on non-susceptible hosts (Figure 2D). The average number of mutations remained high after selection, with 5.28±0.84 mutations for 6mer motifs and 8.87±1.1 mutations for 10mer motifs with a similar distribution as before selection (Figure S2A). There was no statistical correlation between the number of mutations and activity (Figure S2B). The average number of mutations remained high as activity increased for 6mer motifs, while 10mer motifs saw a slight shift towards fewer mutations as activity improved (Figure 2E). Overall, this indicated that activity was not restricted to motifs similar to the wildtype sequence.

Active variants were distributed throughout the tip domain, with most active variants containing motifs in the C β-sheet and BC exterior loop regions (Figure 2F). Variant with improved activity on non-susceptible hosts were more distributed throughout the tip domain (Figure 3A, Figure S2C). The large number of viable variants with motifs in the C β-sheet and BC exterior loop contrasts the initial distribution of motifs, where most motifs were in the DE exterior loop and E β-sheet region, indicating the more BC-C region may be more tolerant of significant alterations. There are several possible reasons for greater preference in this region. First, the native sequence in the local region has no charged residues and four glycine residues across a short area, indicating the native region may be less reactive and more tolerant to larger disruptions. Additionally, this region faces ‘outwards’ away from possibly destabilizing interactions with other monomers of the trimer RBP and may be well positioned to interact with host receptors or other host surface proteins to facilitate successful infections [34].

**Figure 3.**
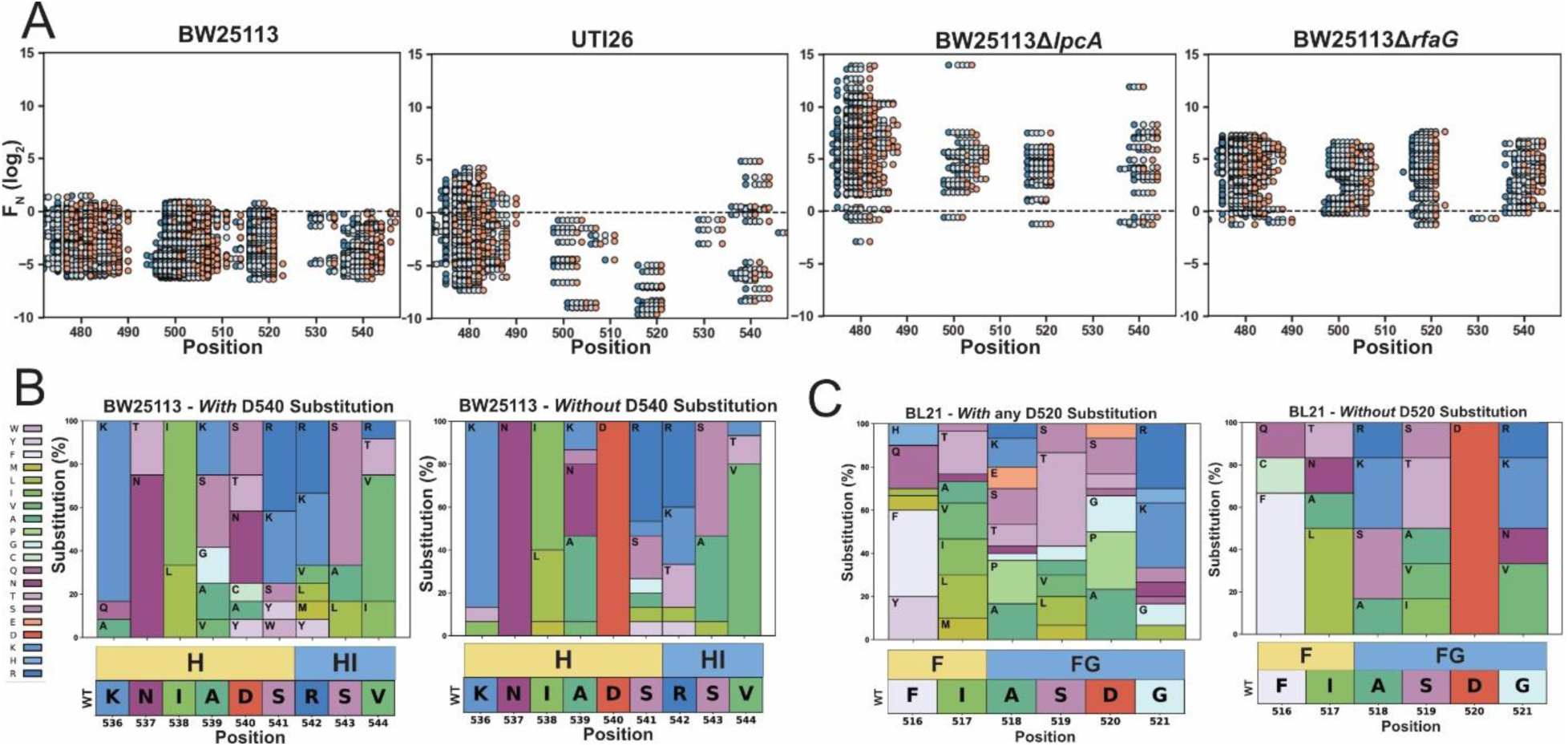
Motifs are distributed across the tip domain. **(A)** The F_N_ score and position in gp17 of detectable phage variants across different hosts. Each point displays a substitution for a motif, with the start of the motif shaded dark blue and the final substitution light red. **(B-C)** Substitution frequency plots for (B) positions 536 through 544 for variants on BW25113 with a substitution at D540 (left) or without a substitution at D540 (right) and (C) positions 516 through 521 for variants on BL21 with a substitution at D520 (left) or without a substitution at D520 (right). The wildtype sequence and topology are shown bottom, and amino acids are colored based on similarity of physicochemical properties as shown left in (B).

The motifs allowed us to examine how multiple substitutions impacted positions essential to activity in susceptible hosts. We analyzed the substitution preference surrounding two conserved regions for variants expected to have activity (F_N_ > -3 for susceptible strains, or at least 1/8^th^ wildtype activity) on BL21 and BW25113. We had previously noted that D520 was highly conserved in T7 phage on *E. coli* BL21 and D540 was highly conserved for BW25113. However, many active motifs contained a substitution at either position. Examining variants with and without a D540 substitution on BW25113 revealed similar substitution profiles with positively charged or uncharged polar substitutions bracketing the position, indicating the negatively charged residue could be fully replaced in this strain (Figure 3B). For BL21, variants that replaced D520 had enriched proline substitutions at A518 and D520, which may cause structural perturbations that could influence receptor recognition (Figure 3C). Some variants also substituted glutamate at this position, a similar negatively charged residue that could recover receptor binding, indicating a negative charge in this region is likely still optimal for maintaining phage activity.

Overall, several important insights emerged from the screen. 24.5% of variants with a sequence motif demonstrated activity, constituting a substantial fraction of the input library. For comparison, our prior screen of single amino acid mutations produced less than 40% of functional phage variants on *E. coli* 10G, while variants in this library contain approximately 5 (for 6mer) or 9 (for 10mer) mutations per sequence after selection. The shift in preference for the BC-C region indicates that position and composition are important for engineering and improving activity in the phage, and motifs that can improve activity non-susceptible strains and replace previously essential amino acids were abundant in the library. Altogether these results suggest that Meta-SIFT unearthed substantially diverse sequence motifs that are tolerated by the RBP and able to confer a fitness benefit to the phage.

### Hierarchical Clustering Reveals Patterns in Motif Selection

To understand how motif preferences governed phage specificity and influenced host range, we examined the activity of the phage library across each bacterial host, merging 6mer and 10mer motifs together. The specificity profile of the phage library was diverse, with host specificity ranging from variants active only on a single host to broadly active variants with activity on up to 19 hosts or conditions in the evaluated panel (Figure 4A).

**Figure 4.**
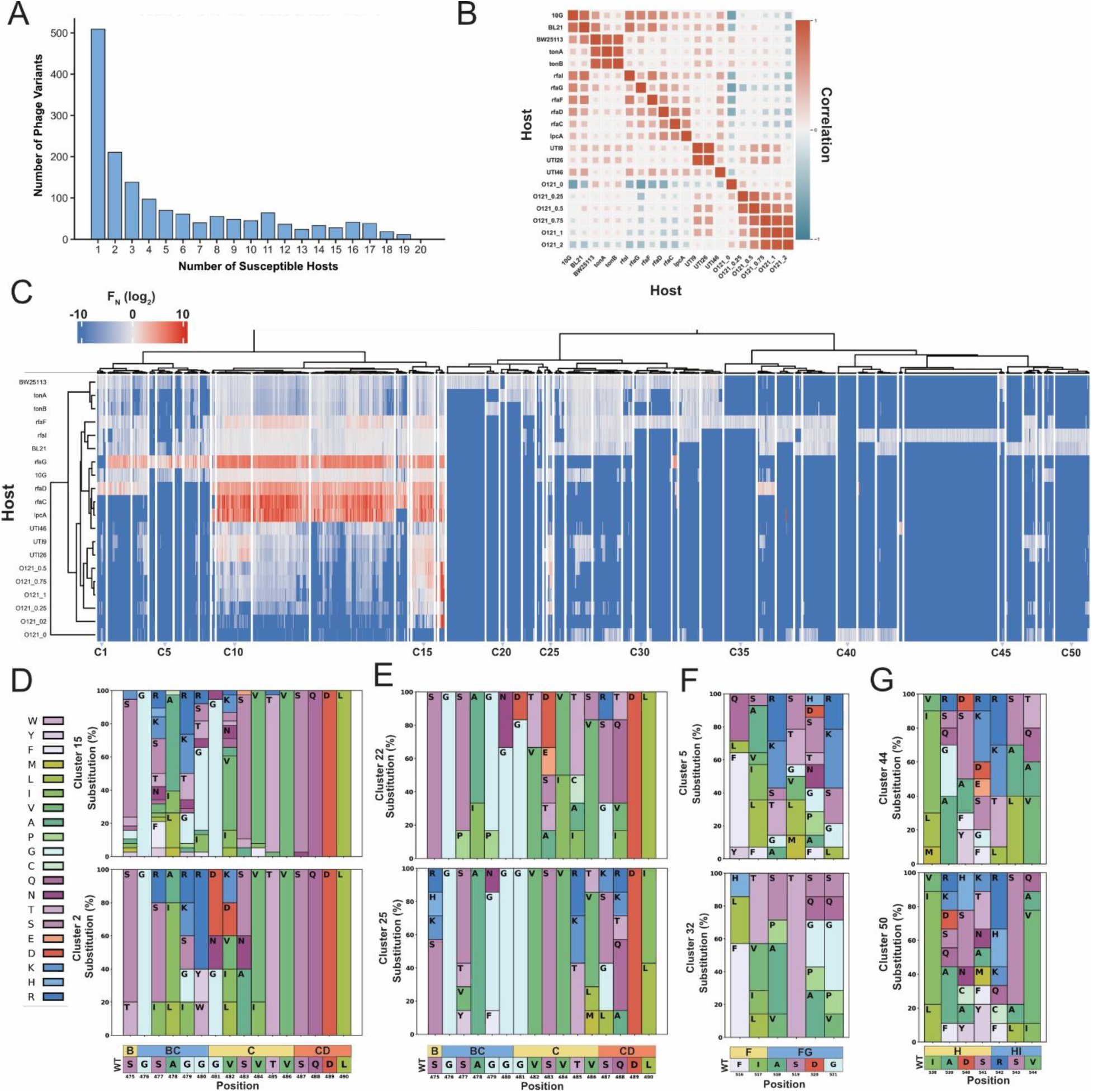
Hierarchical clustering reveals patterns in motif selection. **(A)** The total number of phage variants (inclusive of all 6mer and 10mer motifs) and the associated number of expected susceptible hosts (F_N_ > -3) for each variant. **(B)** Spearman correlation of phage library activity between each pair of hosts, shown as a gradient from red (1, perfectly correlated) to white (0, no correlation), to blue (-1, perfectly negatively correlated). **(C)** Hierarchical clustering of normalized functional scores (log_2_ F_N_, blue to red gradient, wildtype F_N_ = 0) for active phage variants on twenty *E. coli* hosts or conditions (listed left, identified by the strain name, LPS deletion, or for O121 the relevant salt concentration). F_N_ clustered is the average of two biological replicates, spliced into 50 clusters. **(D-G)** Substitution frequency plots comparing substitution frequency for (D) clusters 15 and 2 at positions 475 through 490, (E) clusters 22 and 25 at positions 475 through 490, (F) clusters 5 and 32 at positions 516 through 521, and (G) clusters 44 and 50 at positions 538 to 544. The wildtype sequence and topology are shown bottom, and amino acids are colored based on similarity of physicochemical properties as shown left in (D).

Correlating the F_N_ scores of variants across all hosts revealed broad groupings of host preference (Figure 4B). Phage activity on susceptible hosts 10G and BL21 was more highly correlated to each other and some BW25113 LPS truncations than to wildtype BW25113. BW25113Δ*tonA* and BW25113Δ*tonB* correlated well with BW25113, as these deletions are not expected to influence LPS receptor recognition. LPS deletions resulting in similar truncations had higher correlation. Uropathogenic strains UTI9 and UTI26 highly correlated with each other and 0.5% to 1% NaCl concentrations on *E. coli* O121. The activity profile for the library on *E. coli* O121 negatively correlated with most other hosts, indicating a tradeoff in activity, especially at 0% and 2% NaCl.

To reveal sequence determinants driving host specificity we performed unsupervised hierarchical clustering of the library based on their activity on each bacterial host. Correlation-based agglomerative hierarchical clustering (Spearman distance, Ward’s clustering metric) was used due to its tolerance of outliers and ability to identify smaller clusters, making it ideal for granular classification of host specificities. We grouped variants with similar activity profiles into 50 clusters based on a measurement of optimal cluster density and ease of visualizing trends in specificity (C1-C21, Figure 4C and Figure S3A). The number of clusters used to segregate the dendrogram does not change the relationships between each host and phage variant (Figure S3B-C).

Examination of the hierarchical clustering revealed how both locations where motifs were incorporated on the tip domain and substitution patterns drove host specificity. For example, cluster 15 (38 members) contained variants with broad activity on susceptible strains, uropathogenic *E. coli*, BW25113 LPS mutants, and on *E. coli* O121 in most conditions besides 0% salt. Variants in this cluster typically contained positively charged and polar uncharged substitutions in the BC exterior loop (Figure 4D, Top). In comparison, cluster 2 (40 members) lost activity on O121, uropathogenic strains, and some BW25113 variants with LPS truncations. This cluster notably incorporated negatively charged substitutions into positions in the C β-strand, including mutations at position G481 that were usually not mutated in cluster 15 (Figure 4D, Bottom).

Other clusters contained mutations farther along the C β-strand into the CD interior loop. Cluster 22 (20 members) predominately contained negatively charged substitutions in the C β-strand and occasional proline substitutions in the BC exterior loop (Figure 4E Top). Variants in this cluster generally only had activity on BW25113, BW25113Δ*tonA*, and BW25113Δ*tonB*. In contrast, cluster 25 (8 members), contained variants with a similar activity profile to broadly active cluster 15 except without activity on hosts with truncated LPS. These variants never retained a positively charged substitution in the BC exterior loop, but instead in the surrounding β-sheets, with these substitutions extending into the CD interior loop on the opposite side of the tip domain, indicating these regions are relevant for both host specificity and efficacy but are exclusionary to hosts with the LPS truncations (Figure 4E Bottom).

Substitutions driving specificity were more dramatic for other clusters. Cluster 5 (27 members) contained positively charged and polar uncharged substitutions into the FG exterior loop, usually replacing negatively charged D520. This cluster had activity on 10G, BL21, and BW25113Δ*rfaG* while retaining only weak activity on BW25113 (Figure 4F Top). In contrast, cluster 32 (8 members) lost activity on 10G and BL21 while improving activity on BW25113 and BW25113Δ*tonA*, BW25113Δ*tonB*, and BW25113Δ*rfaF*. Variants in this cluster universally replace D520 and otherwise tended to incorporate polar uncharged or proline residues into the exterior loop, suggesting a change in secondary structure may be responsible for host specificity (Figure 4F Bottom). Other clusters showed how small compositional changes could refine specificity. Cluster 44 (164 members) was characterized with activity only on BW25113Δ*rfaI*, while cluster 50 (60 members) generally had activity on *BW25113ΔrfaI* and BL21. The substitution profile across the H β-strand and HI exterior loop was similar for both clusters, with a different preference for the location of negatively charged substitutions, either at D540 or S541 for cluster 44 or A539 for cluster 50.

These results show that both the location and the composition are key drivers of phage host specificity. Interactions between motifs with many substitutions are complex, but broad trends can be gleaned through analysis of patterns of substitution to reveal novel combinations that can contract or expand host range. Notably, we found no variants that were functional on every host, supporting the idea that loss of activity in one condition may be required to improve activity in others.

### Meta-SIFT identifies motifs highly active on foodborne pathogen *E. coli* O121

*E. coli* (STEC) O121 is one of the six non-O157 serogroups of *E. coli* commonly responsible for foodborne outbreaks [35–38]. This bacterium has not been previously reported as susceptible to bacteriophage T7. Here we demonstrate wildtype T7 activity is, in fact, contingent on a salt deficit (Figure 1C). Interestingly, this indicates the O-antigen does not sterically prevent the phage from binding to its receptor as has been proposed for other phages [39]. Phage variants active on this host are especially notable due to the prominence of the bacteria as a foodborne pathogen.

In higher salt conditions this host is completely resistant to both wildtype phage and a DMS library saturating all single amino acid substitutions in the tip domain (Figure 2B). This contrasts non-susceptible hosts like BW25113Δ*rfaC*, where passage with high concentrations of natural phage can reveal rare mutants that form plaques and passage with the DMS library reveals patterns of single amino acid substitutions that can rescue activity. Engineering an active phage on this host is thus a novel challenge, as a combinatorial solution to improve activity is too far in sequence space from the wildtype sequence to be present in the natural diversity of the phage population.

Examination of successful variants across different salt conditions revealed distinct patterns of substitutions. Every enriched variant in 2% salt had substitutions in either the β-sheet at positions S475 or in the C β-sheet at positions G481 or V482 as well as in the BC exterior loop (Figure 5A), emphasizing the importance of this region. In conditions of lower salt concentration some active variants extended substitutions up the C β-sheet into the CD interior loop of the tip domain, suggesting these regions are amenable to substitution and engineering. Conditions with salt frequently enriched variants with positively charged substitutions at one or more of S475, S477, G479 and V482, while variants with these substitutions were depleted without salt at all positions besides V482 (Figure 5B, all variants with F_N_ > -3). Variants active without salt were instead enriched in more negatively charged residues at multiple positions. Polar, uncharged substitutions were also enriched, with less frequency in 2% salt, and across all conditions G481 remained either unchanged or replaced with a disruptive negatively charged aspartic acid or polar, uncharged asparagine. The variants most enriched across different conditions (Figure 5C, displaying 2% NaCl F_N_ > 3, 0% NaCl F_N_ > 0) showed a similar pattern of substitution to these trends.

**Figure 5.**
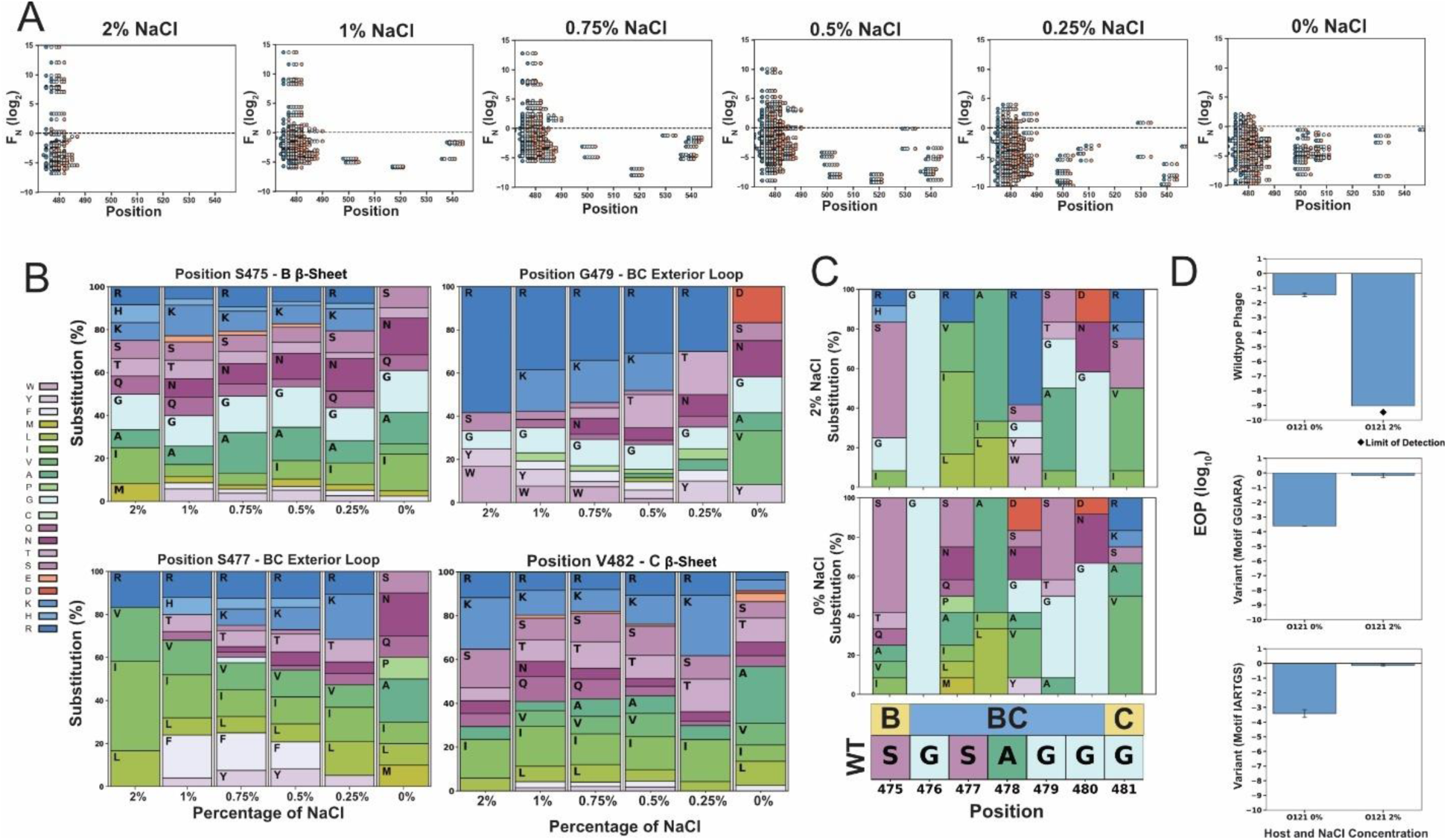
Meta-SIFT identifies active motifs on foodborne pathogen *E. coli* O121. **(A)** The F_N_ score and position in gp17 of detectable phage variants across different salt conditions (2% to 0%). Each point displays a substitution for a motif, with the start of the motif shaded dark blue and the final substitution light red. **(B)** Substitution frequency plots at positions S475 (top left), S477 (bottom left), G479 (top right), and V482 (bottom right) in different salt conditions for variants in each condition where F_N_ > -3. Amino acids are colored based on similarity of physicochemical properties as shown left. **(C)** Substitution frequency plots from positions 475 to 481 for 2% (top, variants with an F_N_ > 3) and 0% (bottom, variants with an F_N_ > 0). The wildtype sequence and topology are shown bottom, amino acids are colored by physicochemical properties as shown in (B). **(D)** Efficiency of Plating (EOP, log_10_) on *E. coli* O121 for wildtype T7 (top) for reference, T7 with motif GGIARA from position 475 to 480 (middle), and T7 variant with motif IARTGS from positions 477 to 482 (bottom). EOP normalized to *E. coli* 10G a wildtype gp17 helper plasmid. Shown as the average of three replicates +/- SD.

To confirm our screen identified variants that truly had activity on *E. coli* O121 in high salt conditions, we directly synthesized (see methods) high scoring variants with motifs GGIARA (Variant 1 - S475G, S477I, G479R, G480A) from position 475 to 480 and IARTGS (Variant 2 - S477I, G479R, G480T, V482S) from positions 477 to 482 and determined plaque efficiency through efficiency of plating experiments. Indeed, engineering variants with these motifs enabled phage variants to productively plaque on *E. coli* O121 in 2% salt, indicating our approach was successful (Figure 5D). These phages also demonstrate lytic capability in liquid culture (Figure S4). As expected, these mutations presented a tradeoff with a reduced ability to plaque on O121 without salt compared to the wildtype phage. Overall, these results paint a complex picture for engineering phage activity on *E. coli* O121 across different conditions. Multiple substitutions across loops and β-sheets are required to enable activity, and identifying these variants was only possible using Meta-SIFT given the total lack of activity in higher salt conditions.

## Discussion

In this study, we devised Meta-SIFT as a new approach engineer complex changes in phage sequences by leveraging diverse metagenomic databases to find metagenomic motifs that influence phage activity. Using Meta-SIFT, we isolated 10^4^ 6-mer and 10-mer motif candidates for the T7 RBP that significantly differed from the wildtype sequence. These motifs were sourced from disparate and distantly related structural metagenomic sequences, allowing us to overcome a lack of sequence conservation among phages to find functionally important motifs. By screening phage variants with these motifs, we revealed thousands of T7 phage variants with novel host specificity and elucidated how motifs drove changes activity in the RBP.

We demonstrate that leveraging DMS results was a successful strategy for correctly seeding scores to find motifs that broadly influenced phage activity and altered host specificity, even if the motifs differ substantially in sequence identity [2]. This limits the method to those phages characterized using high throughput methods like DMS. However, high throughput approaches to characterizing phages are increasingly available, and as phages are developed as therapeutic options more data accumulates for relevant phages that could be used to inform Meta-SIFT.

Using Meta-SIFT, we engineered T7 bacteriophage, usually considered a simple laboratory phage unsuitable for clinically relevant applications, to become highly efficacious on *E. coli* O121, a prominent foodborne pathogen. Historically, phage engineering has heavily relied on natural diversity or targeted or untargeted randomization to tailor phage to different conditions, but these approaches cannot effectively address the enormous combinatorial size and potential of the phage genome. The mutational space in any phage vastly exceeds our ability to drive phages by evolutionary passage. Meta-SIFT exemplifies the utility of refining this combinatorial space to isolate the most relevant regions and substitutions to engineer. For example, an alternative approach is defining small regions to attempt to fully randomize very short regions, as has been performed for exterior loops in T3 phage, or random mutagenesis [23]. The first approach is viable only for extremely short regions due to rapidly expanding combinatorial space and the second approach cannot explore the space to any effective degree. Indeed, the 8 amino-acid stretch that was enabling for *E. coli* O121 comprises a total combinatorial space of 2.56×10^10^ possible sequences, far exceeding the number of sequences that can evaluated effectively without a refining approach like Meta-SIFT.

Both the position and composition of motifs were important drivers of phage activity. The BC exterior loop and C β-strand were overall more tolerant of substitution, although motifs inserted at other positions could drive host specificity. Novel combinations of substitutions appear to allow for substantial changes to the local architecture of the tip domain that enable key residues to contact the host receptor. Unbiased libraries of phages with Meta-SIFT-identified motifs can reveal potent combinations of mutations that can tailor host range to specific hosts. These results further emphasize the need for an unbiased approach to library creation like ORACLE, given the poor correlation of library activity between different hosts. Notably, we found no variants that are functional across every host. These results support the idea of fundamental tradeoffs in host specificity and suggest that engineering a phage to target multiple hosts across different conditions is best done by tailoring a population of that phage, rather than relying on one sequence. Altogether these results improve our understanding of the sequence-function landscape of the T7 RBP and identify motifs that can drive highly specific changes in host activity.

Vast troves of phage metagenomic data are becoming increasingly available from diverse biomes. The enormous breadth of phage diversity makes querying these databases for relevant sequences a challenging prospect. Meta-SIFT is designed as a process that can tap into these complex datasets to identify relevant motifs that drive phage activity. This process is not limited to the RBP and can feasibly be used in any highly characterized region to seed and score potential motifs, including other structural proteins, lysins or holins. Meta-SIFT allows us to move beyond sequence similarity to deeply mine metagenomic databases for relevant sequence motifs. We expect Meta-SIFT can be improved with more expansive datasets, improving the accuracy of predicted motifs. Through leveraging these metagenomic motifs we envision Meta-SIFT as a platform to improve our understanding of sequence-function relationships and engineer phages.

## Acknowledgements

We thank Dr. Rodney Welch for UTI strains and Dr. Douglas Weibel for BW25113 deletion mutant strains. This work was supported by National Institute of Allergy and Infectious Diseases (NIAID) grant R21AI156785 (to S.R.), by funding from the National Institute of General Medical Sciences of the National Institutes of Health grant R35GM143024 (to K.A) and by a Wisconsin Distinguished Graduate Fellowship Award from the University of Wisconsin-Madison and the William H. Peterson Fellowship Award from the University of Wisconsin-Madison Department of Bacteriology (to K.K.).

## Contributions

P.H.: Conceptualization, Data curation, Software, Formal analysis, Investigation, Visualization, Methodology, Writing - original draft, Writing - review and editing

K.K.: Conceptualization, Data curation, Software, Formal analysis, Investigation, Methodology, Writing - review and editing

A.M.: Software

K.N.: Software

K.A.: Conceptualization, Resources, Supervision, Funding acquisition, Project administration, Writing - review and editing

S.R.: Conceptualization, Resources, Supervision, Funding acquisition, Project administration, Writing - review and editing

## Supplementary Figures

**Figure S1.**
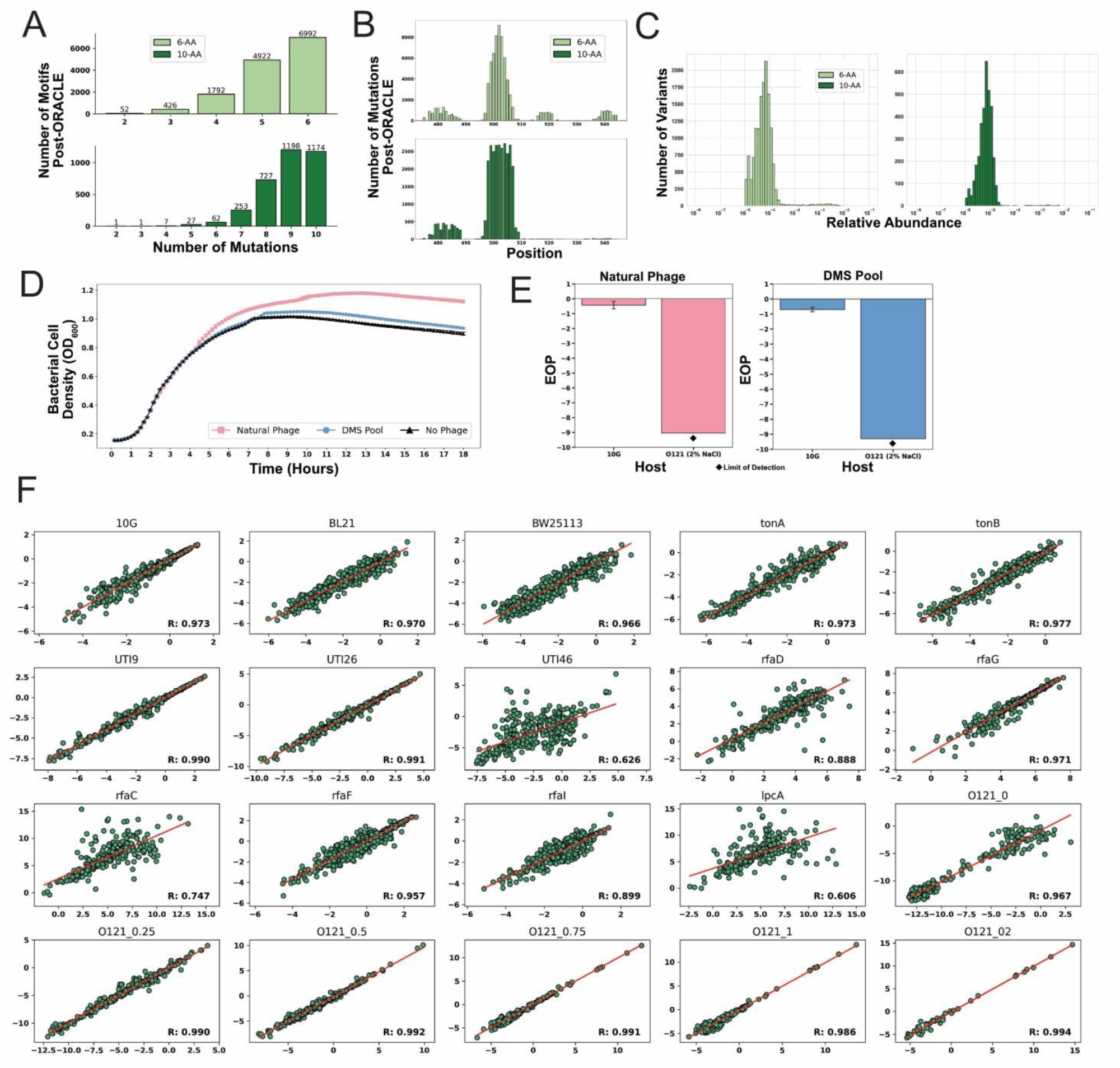
**(A)** Number of mutations per sourced motif for 6-AA (light green, top) or 10-AA (dark green, bottom) motifs in variants in the final expressed library following ORACLE. **(B)** The number of and relative position on the RBP of each mutation for 6mer (top, light green) and 10mer (bottom, dark green) motifs derived from the metagenomic dataset in variants in the final expressed library following ORACLE. **(C)** Abundance of variants in the expressed pool of phages after ORACLE for 6-AA motifs (light green, left) or 10-AA motifs (dark green, right). **(D)** Growth curve on *E. coli* O121 after adding an MOI of 10 of natural wildtype T7 (pink) or a DMS library that includes the tip domain (blue). Control with no phage shown in black. Shown as the average of three replicates +/- SD. **(E)** Efficiency of plating (EOP, log_10_) for natural phage (left, pink) or DMS library (right, blue) on *E. coli* 10G and *E. coli* O121. EOP normalized to *E. coli* 10G a wildtype gp17 helper plasmid. Shown as the average of three replicates +/- SD. **(F)** Correlation plot of F_N_ scores across biological duplicates for each evaluated host. *E. coli* BW25113 knockouts are labeled with the knocked-out gene, variable NaCl conditions for *E. coli* O121 are shown as O121 and the salt concentration.

**Figure S2.**
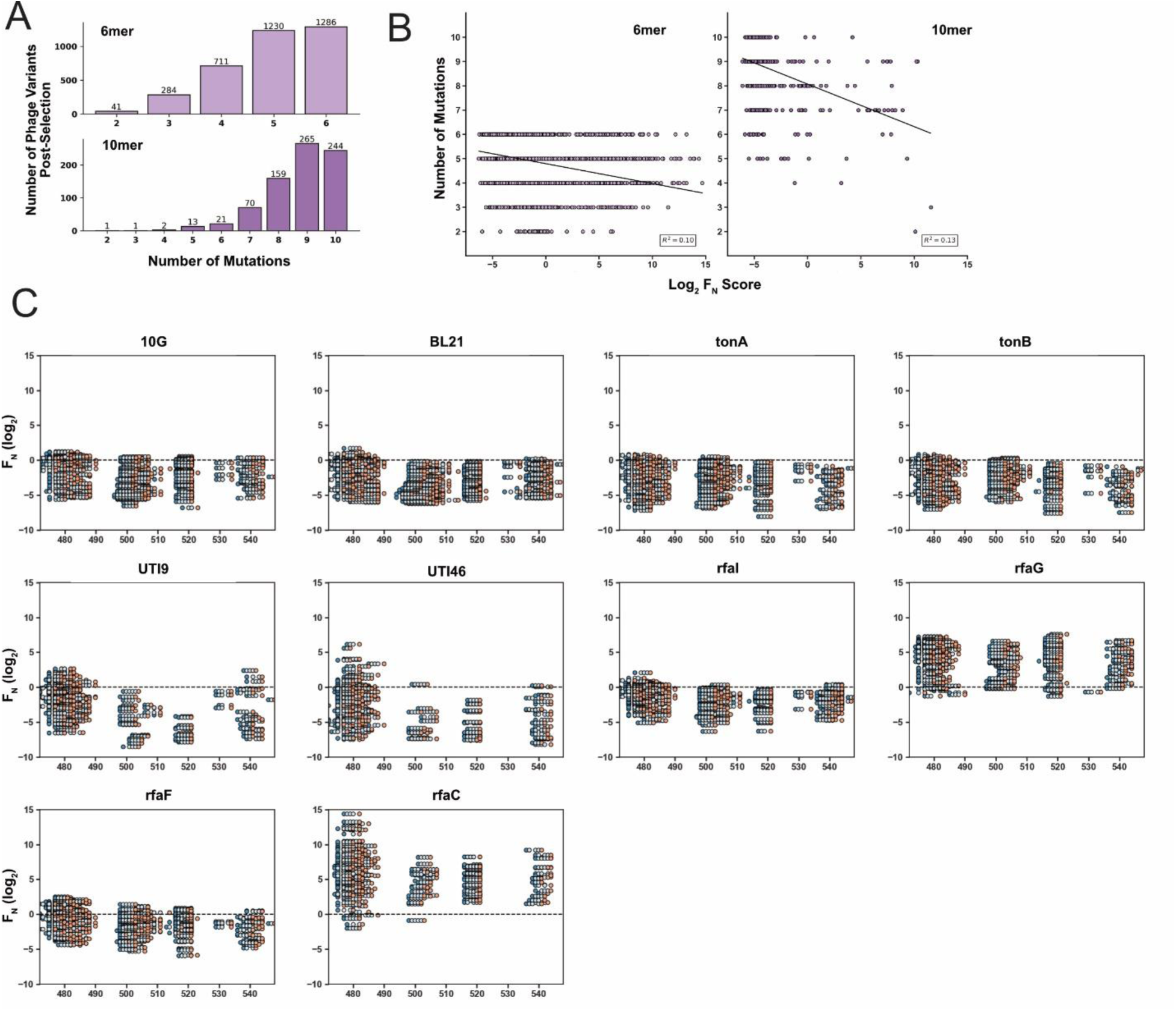
**(A)** Number of mutations per sourced motif for 6-AA (light purple, top) or 10-AA (dark purple, bottom) motifs in variants after host selection. All results are aggregated for all hosts. **(B)** Correlation between the number of mutations in a motif and associated F_N_ score for 6-AA motifs (left, light purple) and 10-AA motifs (right, dark purple) **(C)** The F_N_ score and position in gp17 of detectable phage variants across all remaining hosts. Each point displays a substitution for a motif, with the start of the motif shaded dark blue and the final substitution light red.

**Figure S3.**
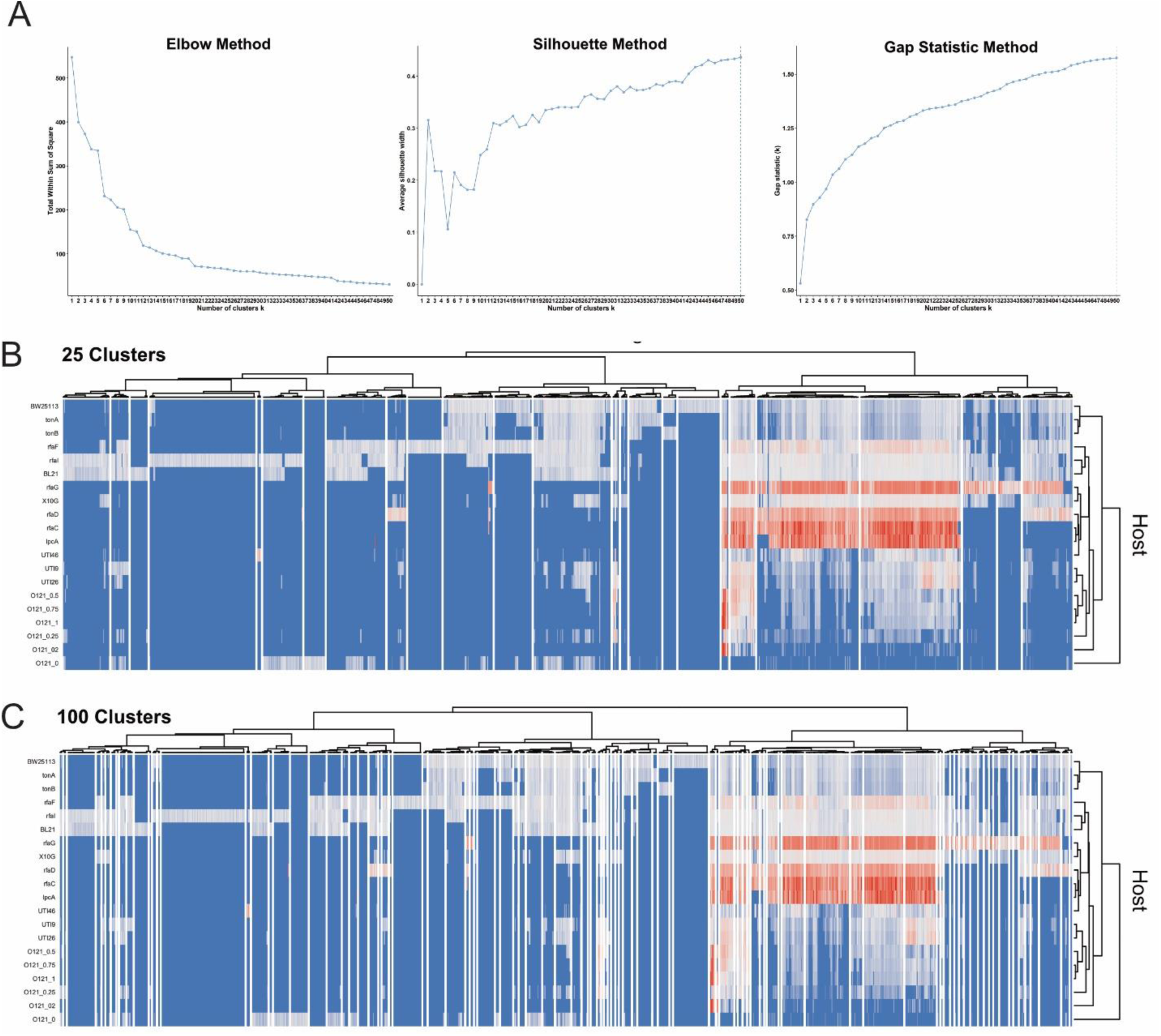
**(A)** Suggested number of clusters using the Elbow (left), silhouette (middle) and gap statistic (right) methods. **(B-C)** Hierarchical clustering of normalized functional scores (log_2_ F_N_, blue to red gradient, wildtype F_N_ = 0) for active phage variants on twenty *E. coli* hosts or conditions (listed left, identified by the strain name, LPS deletion, or for O121 the relevant salt concentration). F_N_ clustered is the average of two biological replicates, shown split into (B) 25 or (C) 100 clusters.

**Figure S4.**
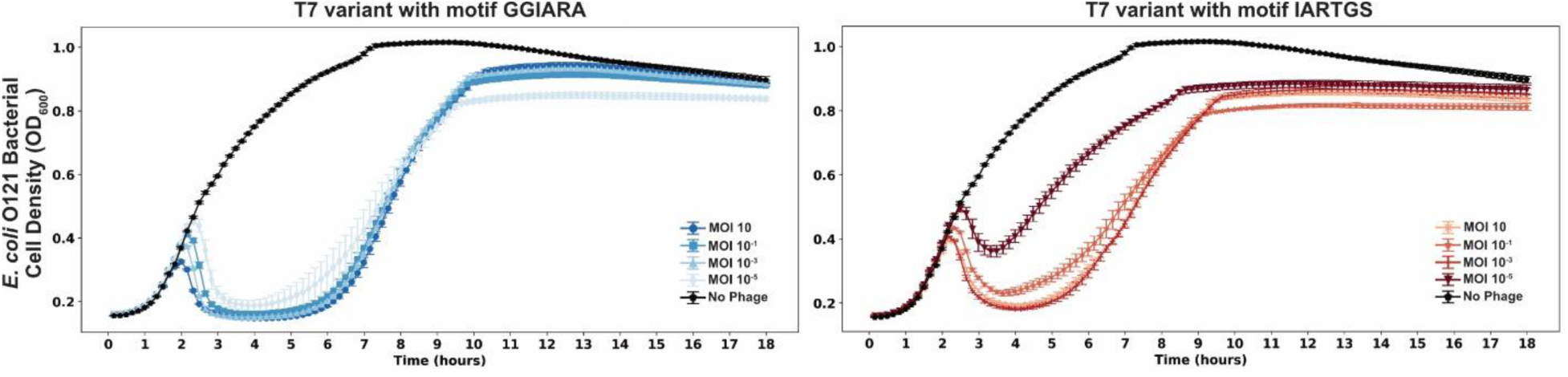
Growth curve on *E. coli* O121 after adding different MOIs of T7 variant with motif GGIARA (left, blue gradient) or T7 variant with motif IARTGS (right, red gradient). Control with no phage shown in black. Shown as the average of three replicates +/- SD.

## Methods

### Microbes and Culture Conditions

T7 bacteriophage was obtained from ATCC (BAA-1025-B2). *Escherichia coli* BL21 is a lab stock, *E. coli* 10G is a highly competent DH10B derivative [40] originally obtained from Lucigen (60107-1). *E. coli* BW25113 and knockout strains were obtained from Doug Weibel (University of Wisconsin, Madison) and are derived from the Keio collection [41]. UTI strains were obtained from Rod Welch (University of Wisconsin, Madison) and originates from a UTI collection [42]. *E. coli* O121 was obtained from ATCC (BAA-2190). All bacterial hosts are grown in and plated on Lb media (1% Tryptone, 0.5% Yeast Extract, 1% NaCl in dH2O, plates additionally contain 1.5% agar, while top agar contains only 0.5% agar). For experiments with variable salt conditions Lb is made without salt and salt added as needed. Lb media was used for all experimentation and was used to recover host and phages after transformation. Kanamycin (50 μg/ml final concentration, marker for pHT7Helper1) and spectinomycin (115 μg/ml final concentration, marker for pHRec2, pHRec1-Motif and pHCas9 and derivatives) was added as needed. All incubations of bacterial cultures were performed at 37°C, with liquid cultures shaking at 200-250 rpm unless otherwise specified. Bacterial hosts were streaked on appropriate Lb plates and stored at 4°C.

T7 bacteriophage was propagated using *E. coli* BL21 after initial receipt from ATCC and then as described on various hosts in methods. All phage experiments were performing using Lb and culture conditions as described for bacterial hosts. Phages were stored in Lb at 4°C.

For long term storage all microbes were stored as liquid samples at -80°C in 10% glycerol, 90% relevant media.

SOC (2% tryptone, 0.5% yeast extract, 0.2% 5M NaCl, 0.25% 1M KCL, 1% 1M MgCl_2_, 1% 1M MgSO_4_, 2% 1M glucose in dH2O) was used to make competent cells.

### General Cloning Methods

PCR was performed using KAPA HiFi (Roche KK2101) for all experiments. Golden Gate assembly was performed using New England Biosciences (NEB) Golden Gate Assembly Kit (BsaI-HFv2, E1601L). Restriction enzymes were purchased from NEB. DNA purification was performed using EZNA Cycle Pure Kits (Omega Bio-tek D6492-01) using the centrifugation protocol. Gibson assembly was performed according to the Gibson Assembly Protocol (NEB E5510) but Gibson Assembly Master Mix was made in lab (final concentration 100 mM Tris-HCL pH 7.5, 20 mM MgCl_2_, 0.2 mM dATP, 0.2 mM dCTP, 0.2 mM dGTP, 10 mM dTT, 5% PEG-8000, 1 mM NAD^+^, 4 U/ml T5 exonuclease, 4 U/μl Taq DNA Ligase, 25 U/mL Phusion polymerase). Acceptor phages were made using NEB NEBuilder HiFi DNA Assembly Gibson mix. All cloning was performed according to manufacturer documentation except where noted in methods.

PCR reactions use 1 μl of ∼1 ng/μl plasmid or ∼0.1 ng/μl DNA fragment as template for relevant reactions. PCR reactions using phage as template use 1 μl of undiluted phage stock (genomic extraction was unnecessary) and have an extended 5-minute 95°C denaturation step.

DpnI digest was performed on all PCR that used plasmid as template. Digestion was performed directly on PCR product immediately before purification by combining 1.5 μl DpnI (30 units), 15 μl 10x CutSmart Buffer, 98 ul PCR product, and 35.5 ul dH2O, incubating at 37°C for 2 hours and 30 minutes then heat inactivating at 80C for 20 minutes.

Electroporation of plasmids was performed using a Bio-rad MicroPulser (165-2100), Ec2 setting (2 mm cuvette, 2.5 kV, 1 pulse) using 50 μl competent cells and 100 ng of plasmid or 20 ul of golden gate reaction for transformation. Electroporated cells were immediately recovered with 950 μl Lb, then incubated at 37°C for 1 to 1.5 hours and plated or grown in relevant media.

*E. coli* 10G competent cells were made by adding 8 mL overnight 10G cells to 192 mL SOC (with antibiotics as necessary) and incubating at 21°C and 200 rpm until ∼OD_600_ of 0.4 as determined using an Agilent Cary 60 UV-Vis Spectrometer using manufacturer documentation (actual incubation time varies based on antibiotic, typically overnight). Cells are centrifuged at 4°C, 1000g for 20 minutes, the supernatant is discarded, and cells are resuspended in 50 mL 10% glycerol. Centrifugation and washing are repeated three times, then cells are centrifuged at 2000g for 20 minutes then resuspended in a final volume of ∼1 mL 10% glycerol and are aliquoted and stored at -80°C.

### Plasmid Cloning and Descriptions

<u>pHT7Helper1</u> is used during optimized recombination and during accumulation in ORACLE to prevent library bias and depletion of variants that grow poorly on *E. coli* 10G. There are no changes to this plasmid from its initial construction as documented in our previous DMS assay, plasmid details are provided here for completeness [2]. This plasmid contains a pBR backbone, kanamycin resistance cassette, mCherry, and the T7 receptor binding protein *gp17*. Both mCherry and *gp17* are under constitutive expression. *Gp17* was combined with promoter apFAB47 [43] using SOE and the plasmid assembled by Gibson assembly. There is a single nucleotide deletion in the promoter that has no effect on plaque recovery for phages that require *gp17* to plaque.

<u>pHRec2</u> is used as template to generate the variant library containing metagenomic motifs, referred to as pHRec2-Motif and used during optimized recombination in ORACLE. This plasmid has been updated since its initial construction in our previous DMS assay, where it was called pHRec1. As before this plasmid contains an SC101 backbone, Cre recombinase, a spectinomycin resistance cassette, and the T7 tail fiber *gp17* flanked by Cre lox66 sites with an m2 spacer, a 3’ pad region and lox71 sites with a wt spacer [44]. Cre recombinase is under constitutive expression. This plasmid was altered using sequential PCR and Gibson assembly to include a larger constant region and random, 20 bp barcode between the end of *gp17* and the Cre-lox sites. The total region inserted was 105 bp. This barcode was added to facilitate smaller, rather less expensive NGS reads but was not used in this experimental series.

<u>pHCas9-3</u> is used during Accumulation in ORACLE. There are no changes to this plasmid from its initial construction as documented in our previous DMS assay, plasmid details are provided here for completeness [2]. This plasmid contains an SC101 backbone, a spectinomycin resistance cassette and cas9 cassette with a previously cloned gRNA targeting the T7 acceptor phage [45]. pHCas9 was created with Gibson assembly, while pHCas9-3 was assembled by phosphorylation and annealing gRNA oligos (100 uM forward and reverse oligo, 5 μl T4 Ligase buffer, 1 μl T4 PNK, to 50 μl dH2O, incubate at 37°C for 1 hour, 96C for 6 minutes then 0.1C/s temperature reduction to 23C), then Golden Gate cloning (1 μl annealed oligo, 75 ng pHCas9, 2 μl T4 DNA Ligase Buffer, 1 μl Golden Gate Enzyme Mix, dH20 to 20 ul. Incubation at 37°C for 1 hour then 60C for 5 minutes, followed by direct transformation of 1 ul, plated on Lb with spectinomycin). All plasmid backbones and gene fragments are lab stocks.

### General Bacteria and Phage Methods

Bacterial concentrations were determined by serial dilution of bacterial culture (1:10 or 1:100 dilutions made to 1 mL in 1.5 microcentrifuge tubes in Lb) and subsequent plating and bead spreading of 100 μl of a countable dilution (targeting 50 colony forming units) on Lb plates. Plates were incubated overnight and counted the next morning. Typically, two to three dilution series were performed for each host to initially establish concentration at different OD_600_ and subsequent concentrations were confirmed with a single dilution series for later experiments.

Stationary phase cultures are created by growing bacteria overnight (totaling ∼20-30 hours of incubation) at 37°C. Cultures are briefly vortexed then used directly. Exponential phase culture consists of stationary culture diluted 1:20 in Lb then incubated at 37°C until an OD_600_ of ∼0.4-0.8 is reached (as determined using an Agilent Cary 60 UV-Vis Spectrometer using manufacturer documentation), typically taking 40 minutes to 1 hour and 20 minutes depending on the strain and antibiotic, after which cultures are briefly vortexed and used directly.

Phage lysate was purified by centrifuging phage lysate at 4g, then filtering supernatant through a 0.22 uM filter. Chloroform was not used and is recommended against.

To establish titer, phage samples were typically serially diluted (1:10 or 1:100 dilutions made to 1 mL in 1.5 microcentrifuge tubes) in Lb to a 10^-8^ dilution for preliminary titering by spot assay. Spot assays were performed by mixing 250 μl of relevant bacterial host in stationary phase with 3.5 mL of 0.5% top agar, briefly vortexing, then plating on Lb plates warmed to 37°C. After plates solidified (typically ∼5 minutes), 1.5 μl of each dilution of phage sample was spotted in series on the plate. Plates were incubated and checked after overnight incubation (∼20-30 hours) to establish a preliminary titer. After a preliminary titer was established, phage samples were serially diluted in triplicate for efficiency of plating (EOP) assays.

EOP assays were performed using whole plates instead of spot plates to avoid inaccurate interpretation of results due to spotting error [46]. To perform the whole plate EOP assay, 200-400 μl of bacterial host in exponential phase was mixed with between 5 to 50 μl of phages from a relevant dilution targeting 50 plaque forming units (PFUs) after overnight incubation. The phage and host mixture was briefly vortexed, briefly centrifuged, then added to 3.5 mL of 0.5% top agar, which was again briefly vortexed and immediately plated on Lb plates warmed to 37°C. After plates solidified (typically ∼5 minutes), plates were inverted and incubated overnight. PFUs were typically counted after overnight incubation (∼20-30 hours) and the total overnight PFU count used to establish titer of the phage sample. PFU totals between 10 and 300 PFU were typically considered acceptable, otherwise plating was repeated for the same dilution series. This was repeated at least in triplicate for each phage sample on each relevant host to establish phage titer.

EOP was determined using *E. coli* 10G with pHT7Helper1 as a reference host. EOP values were generated for each of the three dilutions by taking the phage titer on the test host divided by the phage titer on the reference host, and this value was subsequently log_10_ transformed. Values are reported as mean ± SD.

MOI was calculated by dividing phage titer by bacterial concentration. MOI for the T7 variant library after the variant gene is expressed was estimated by titering on 10G with pHT7Helper1.

### Position Seed Scoring

Data from our previous DMS assay of the T7 RBP tip domain was used to seed scores for each possible substitution [2]. The maximum F_N_ of each biological replicate for each substitution from susceptible strains 10G, BL21, and BW25113 in our DMS assay was compressed where if F_N_ ≤ 1 compressed F_N_ (CF_N_) = F_N_ and if F_N_ > 1 then CF_N_ = (F_N_ – 1 / max(strain F_N_) -1) + 1. Scores were compressed to reduce the impact of variants that performed exceedingly well, as we wanted to weight scores higher for variants with functionally different scores across strains (i.e. it matters less if a variant does very well on one strain and comparable to wildtype on other strains, it matters more if it does poorly on one strain and comparable to wildtype or better than wildtype on another, as that is more functionally distinct.)

If the difference between CF_N_ on all susceptible strains was less than 0.5, indicating the functional difference between strains was small, motif seed score MFF = CF_N_, else MFF = CF_N_ + 0.5. This served to seed scores higher if there was a functional difference between susceptible strains. Finally max(*⌊*log(F_N_*⌋*) of each biological replicate for each substitution from resistant strains BW25113Δ*rfaG* and BW25113Δ*rfaD* was calculated and added to MFF. This further weighed substitutions higher if they had recovered function on resistant strains. These final values (see Supplementary File 1) were used to seed scores for every possible substitution to curate motifs.

Note as mentioned in the main text this approach presents a notable tradeoff. Increasing the seed score for mutations that changed host range in the DMS dataset predisposes seeded motifs towards sequences that are, by their nature, not expected to have high activity on all bacterial hosts, and this means much of the library is expected to be non-functional. While we reasoned that this tradeoff is worthwhile (and likely necessary) to identify sequences that can drive distinct changes in host range, reducing or eliminating the MFF ‘bonus’ weight from the differential on susceptible strains may reduce the impact of this effect. Variants would be less diverse but more of the library would likely be functional, if such a change is desired.

### Database construction

NCBI databases (release July 2019) were queried for the term “prokaryotic virus” and limited to sequences of at least 3 kb. The resulting sequences were manually curated by removing non-genomic sequences (e.g., containing the terms ‘open reading frame’ or ‘ORF’) and non-viral sequences. Additionally, the IMG/VR v2.1 (October 2018) database was downloaded and limited to sequences originating from human, wastewater, and animal environments according to IMG/VR metadata[27].

For each dataset, open reading frames were predicted using Prodigal v2.6.3 (-p meta) and proteins were dereplicated using CD-HIT v4.7 (-c 1)[47,48]. To obtain proteins of interest, HMM profiles for 373 viral hallmark proteins were selected from VOGDB v94 (vogdb.org) according to viral hallmark proteins designated by VIBRANT v1.2.1[49]. The selected HMM profiles represented annotations containing the terms *tail, capsid, spike, baseplate, sheath, lysin,* and *holin*. Hmmsearch v3.1b (-e 1e-5) was used to query the database proteins to the selected HMM profiles[50]. In total, 34682 and 26335 non-redundant proteins from NCBI and IMG/VR databases, respectively, were selected for final analysis. Source code is available at https://github.com/raman-lab/Meta-SIFT.

### Motif search

A custom motif search tool (*motif_finder_tool.py*) was used to query each of the protein databases separately for 6-mer motifs (-n 6 -c 1 -e 1e-50 -t 0.8) and 10-mer motifs (-n 10 -c 1 -e 1e-5 -t 0.045). The motif search tool scans n-mer windows of input database amino acids against n-mer windows of input matrix scores. The scores per amino acid window are compared to a set cutoff and predetermined constants. The maximum score per amino acid window across all matrix windows is summed to generate an overall score per protein. The overall score and e-value, calculated based on the number of windows searched, are compared to set thresholds. Any protein passing these thresholds is considered as a candidate protein that is similar to the motifs found within the original input matrix. All amino acid windows used in the generation of the overall score for that protein are extracted. To limit the results to a practical search space, 6-mer and 10-mer motifs were filtered to those that were extracted at least 50 and 15 times, respectively, by the motif search tool across all proteins.

### Motif Variant Plasmid Library Preparation

To create the metagenomic motif variant plasmid library, oligos were first designed and ordered from Agilent as a SurePrint Oligonucleotide Library (OLS 230mers, Barcode 33049511001 OLS, internal freezer stock #2). Every oligo contained a single motif that replaced the wildtype region in the tip domain. We used the most frequently found codon for each amino acid in the *gp17* tail fiber to define the codon for each substitution. Oligos contained BsaI sites at each end to facilitate Golden Gate cloning. Two pools of oligos were used for this library. Oligo pools were amplified by PCR using 0.005 pmol total oligo pool as template (∼0.5 ng total DNA) and 15 total cycles to prevent PCR bias, then pools were purified. pHRec2 was used as template in a PCR reaction to create two backbones for each of the three pools, which was then DpnI digested and purified. Backbones are not otherwise pretreated before Golden Gate Assembly. Golden gate assembly was performed using ∼300 ng of relevant pool backbone and a 1x molar ratio for oligos (∼20 ng) and a small ∼100 bp fragment containing constant regions and a 20 bp randomized barcode (∼15 ng), combined with 2 μl 10x T4 DNA ligase buffer, 1 μl NEB Golden Gate Enzyme mix and dH2O to 20 ul. For the 3’ pool this is a 4-part assembly, for the 5’ pool this is a 3-part assembly. These reactions were cycled from 37°C to 16°C over 5 minutes, 60x, then held at 60°C for 5 minutes to complete Golden Gate assembly and held at 4°C overnight. The following day the reaction was held at 60°C for 5 minutes before dialysis in accordance with NEB guidance. Membrane drop dialysis was then performed on each library pool for 90 minutes to enhance transformation efficiency. The entire 20 μl reaction was then transformed into 50 μl competent *E. coli* 10G cells. Drop plates were made at this point (spotting 2.5 μl of dilutions of each library on Lb plates with spectinomycin) and total actual transformed cells were estimated at ∼5×10^5^ CFU/mL for the 5’ pool at ∼1×10^6^ for the 3’ pool. Each 1 mL pool was added to 4 mL Lb with spectinomycin and incubated overnight, then plasmids were purified. Plasmids concentration was determined by nanodrop and pools were then combined at an equimolar ratio to create the final phage variant pool, denoted as pHRec2-Motif. pHRec2-Motif was transformed into *E. coli* 10G with pHT7Helper1 by transforming 50 ng of plasmid into 60 ul of *E. coli* 10G competent cells. Drop plates were made (spotting 2.5 μl of dilutions of each library onto Lb plates with spectinomycin and kanamycin) and total actual transformed cells were estimated at ∼6.7×10^5^ CFU/mL. The 1 mL library was added to 4 mL Lb with spectinomycin and kanamycin and incubated overnight. This was repeated once to ensure >100 coverage of the entire library and all preparations were combined. This host, *E. coli* 10G with pHT7Helper1 and pHRec1-Motif, was the host used for Optimized Recombination during ORACLE.

### Phage Library Preparation Using ORACLE

T7 Acceptor phages were created using Gibson Assembly using NEBuilder HiFi DNA Assembly Reactions. This contrasts our original approach using YAC Assembly – here phage are directly assembled using fragments up to 10 kb in a linear construct without any plasmid backbone, then directly transformed into *E. coli* 10G [2,51]. This method is much faster, cheaper, easier, and more reliable than YAC Assembly and is highly recommended for construct cloning. T7 Accepters are of similar design but with an added region to facilitate barcode sequencing (which was not used in this experiment). Optimized Recombination was performed by adding T7 Acceptor phages (MOI ∼1) to 50 mL exponential phase 10G with pHT7Helper1 and pHRec2-Motif. Cultures were incubated until lysis and phages were purified, then titer was estimated by spot plate at ∼5×10^7^ variants/mL in a total phage population of ∼1×10^10^ PFU/mL. Accumulation was performed by adding ∼MOI of 1 of recombined phages to 50 mL of exponential phase *E. coli* 10G with pHT7Helper1 and pHCas9-3. Cultures were incubated until lysis then phages were purified. Library expression was performed by adding the accumulated library to 5 mL *E. coli* 10G (with no plasmid) at an MOI of ∼1. Cultures are incubated until lysis and phages were purified. This constitutes the final phage variant library with full expression of the variant *gp17* tail fibers. This phage population is directly sequenced to establish the pre-selection library population.

The DMS library evaluated in S1D comprises an unbiased DMS pool made using ORACLE from positions F463 through the end of the tail fiber.

### Motif Library Selection

All motif library selection experiments were performed in the same way. The T7 variant library was added to 5 mL of exponential host at an MOI of ∼2×10^-2^ and the culture was allowed to fully lyse. This MOI was chosen to give a baseline of 2 replication cycles for the phage library. The entire process was repeated in biological duplicate for each host. Phages were then purified and sequenced.

### Deep Sequencing Preparation and Analysis

We used deep sequencing to evaluate phage populations. We first amplified the tip domain by two step PCR, or tailed amplicon sequencing, using KAPA HiFi. Primers for deep sequencing attach to constant regions adjacent to the tip domain, 3’ of the new barcode in pHT7Rec2. Constant regions are also installed in the fixed region of the acceptor phages for the similarly size amplicon so acceptor phages can also be detected. The first PCR reaction adds an internal barcode (used for technical replicates to assay PCR skew), a variable N region (to assist with nucleotide diversity during deep sequencing, this is essential for DMS libraries due to low nucleotide diversity at each position), and the universal Illumina adapter. Four forward and four reverse primers were used in each reaction, each with a variable N count (0, 2, 4, or 8). Primers were mixed at equimolar ratios and total primers used was per recommended primer concentration. PCR was performed using 13 total cycles in the first PCR reaction. The product of this reaction was used directly in the second PCR reaction, which adds an index and the Illumina ‘stem’. This PCR was run for 9 total PCR cycles. The product of this reaction was purified and was used directly for deep sequencing. Each phage population was sampled at least twice using separate internal barcodes, and no PCR reactions were pooled. Total PCR cycles overall for each sample was kept at 22x to avoid PCR skew. All phage samples were deep sequenced using an Illumina Miseq System, 2×250 read length or an Illumina Nextseq 2×150 read length.

Paired-end Illumina sequencing reads were merged with PEAR v0.9.11 using the default software parameters [52]. Phred quality scores (Q scores) were used to compute the total number of expected errors (E) for each merged read and reads exceeding an Emax of 1 were removed [53]. Wildtype, acceptor phages, and each variant were then counted in the deep sequencing output. We correlated read counts for each technical replicate to determine if there was any notable skew from PCR or deep sequencing. Technical PCR replicates correlated extremely well (R≥0.98 for all samples) indicating no relevant PCR skew and were aggregated for each biological replicate. Besides wildtype and acceptor counts, we included only sequences with an entirely correct sequence and known motifs in our analysis, greatly reducing the possibility of deep sequencing error resulting in an incorrect read count for a variant. With this in mind, and to avoid missing low abundance members after selection, we used a low read cutoff of 4. A pseudo-count was not utilized.

To score enrichment for each variant we used a basic functional score (F), averaging results of the three biological replicates where 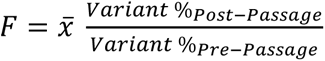. To compare variant performance across hosts we normalized functional score (F_N_) to wildtype, where 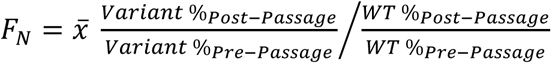.

### Other Statistical Analysis, Data Availability and Source Code

Hierarchical Clustering was performed using the Complex Heatmaps v2.10 package in R [54]. Different distance values were computed using get_dist in R, clustering approaches were calculated using hclust in R, and cophenetic scores were calculated using cophenetic in R. Frequency plots were made in python. P-values for EOP graphs compare plaque capability on the tested host to the reference host for the EOP graph. Counting motif variants was done in Pandas. Relevant statistical details of experiments can be found in the corresponding figure legends or relevant methods section.

## Notes

### Competing Interest Statement

The authors have declared no competing interest.

### Summary of Updates

This manuscript has been significantly revised to include a much larger host panel, including foodborne pathogen E. coli O121 in conditions where the natural phage has no activity. Every figure has had a substantial update to reflect these changes and emphasize the utility of this approach to engineer bacteriophages.

